# Translational control plays an important role in the adaptive heat-shock response of *Streptomyces coelicolor*

**DOI:** 10.1101/223925

**Authors:** Giselda Bucca, Radhika Pothi, Andrew Hesketh, Carla Möller-Levet, David A. Hodgson, Emma E. Laing, Graham R. Stewart, Colin P. Smith

## Abstract

Stress-induced adaptations require multiple levels of regulation in all organisms to repair cellular damage. In the present study we evaluated the genome-wide transcriptional and translational changes following heat stress exposure in the soil-dwelling model actinomycete bacterium, *Streptomyces coelicolor*. The combined analysis revealed an unprecedented level of translational control of gene expression, deduced through polysome profiling, in addition to transcriptional changes. Our data show little correlation between the transcriptome and ‘translatome’; while an obvious downward trend in genome wide transcription was observed, polysome associated transcripts following heat-shock showed an opposite upward trend. A handful of key protein players, including the major molecular chaperones and proteases were highly induced at *both* the transcriptional and translational level following heat-shock, a phenomenon known as ‘potentiation’. Many other transcripts encoding cold-shock proteins, ABC-transporter systems, multiple transcription factors were more highly polysome-associated following heat stress; interestingly, these protein families were not induced at the transcriptional level and therefore were not previously identified as part of the stress response. Thus, stress coping mechanisms at the level of gene expression in this bacterium go well beyond the induction of a relatively small number of molecular chaperones and proteases in order to ensure cellular survival at non-physiological temperatures.

## Introduction

*Streptomyces* are multicellular bacteria highly abundant in the soil and other ecological niches. They are characterized by a complex mycelial life cycle and by their ability to produce a vast array of secondary metabolites, most notably antibiotics. Like unicellular free living organisms they have a versatile and robust metabolism which allows them to adapt to life in different environments and to quickly adjust to environmental fluctuations such as temperature or pH-changes, scarce nutrient or oxygen availability and exposure to antibiotics commonly occurring in the soil environment ^1 2–5^. Their ability to fine tune gene expression at multiple levels (transcriptional, post-transcriptional, translational, post-translational and protein decay) facilitates rapid changes to their proteome in response to extracellular or intracellular signals. Transcriptomic approaches have been used to gain insights into the regulatory changes that govern cellular metabolism at a system-wide level ^6–10^ Recently, polysome and ribosome profiling have been demonstrated as powerful methods for studying changes in the translation of individual mRNAs under different conditions, as well as global effects on the entire actively translated mRNA population or ‘translatome’ ^11,12,13,14,15,16^.

Control of protein synthesis closely correlates with cellular metabolic states ^17^ and translation initiation is commonly considered the rate limiting step in protein synthesis. Consequently, ribosome loading on a transcript can be a robust indicator of translation efficiency which can be monitored via polysome profiling or ribosome profiling ^18,19^ Applications of these approaches have revealed that, in addition to transcriptional induction, post-transcriptional and post-translational processes determine the response to stress in both eukaryotic and prokaryotic organisms ^20,7,15,21^. Moreover, translational control of operonic genes determines the stoichiometry of the different subunits of enzyme complexes in *E. coli* and *S. coelicolor*^22,14, 21^.

In the present study we monitored global changes in transcription and translation following exposure to a sudden increase in temperature (heat stress) in the model streptomycete, *Streptomyces coelicolor*, to reveal the biological processes and metabolic pathways that become activated or repressed under transient stress conditions. This study represents the first step in the identification of key molecular players governing translational control of the adaptive response to cellular stress in this organism. Because environmental stress and morphological development are closely associated in *Streptomyces*, our findings have relevance to both processes ^23^.

## Results and Discussion

### Transcriptional and translational changes following heat-shock

#### Polysome profiling of the heat-shock response

Global transcriptional responses to oxidative and osmotic stress, heat or pH shock, and nutritional shift-down are well documented in *Streptomyces coelicolor*^2,24–26, 27^. How the changes in transcript levels reflect changes in protein levels is currently less well understood, except for the observation that the correlation between the two is generally low or in some instances even divergent, at least in *Streptomyces*^2,6^ and in *E. coli*^9^ To gain insight into the relative contribution of transcriptional and translational regulation to changing the proteome after heat stress, we have undertaken a global comparative transcriptome/translatome approach to monitor gene expression changes in *S. coelicolor* cultures subjected to heat stress.

*S. coelicolor* cultures grown on cellophane-coated SMMS agar were heat-shocked at 42°C for 15 min as previously described (Bucca et al., 1995). In order to compare the relative contribution of transcriptional and translational control to the heat-shock response, samples were taken from five independent cultures both immediately before and after heat-shock. Parallel samples of mycelium were either stabilized in RNA Protect Bacteria Reagent for total RNA extraction, or were treated with chloramphenicol to block translation elongation prior to polysome fractionation and subsequent extraction of the actively translated (ribosome-associated) mRNA populations.

The profiles of the respective polysome sucrose gradients showed differences in profiles where the peaks corresponding to the 30S and 50S subunits and intact 70S monosome were more pronounced in the heat-shocked cultures compared to those in the 30°C cultures (Fig. 1). Strikingly, a fourth peak was also more abundant, following the 70S monosome peak (designated ‘P1’) and is considered to comprise two ribosomes (a ‘disome’).

**Figure 1.**
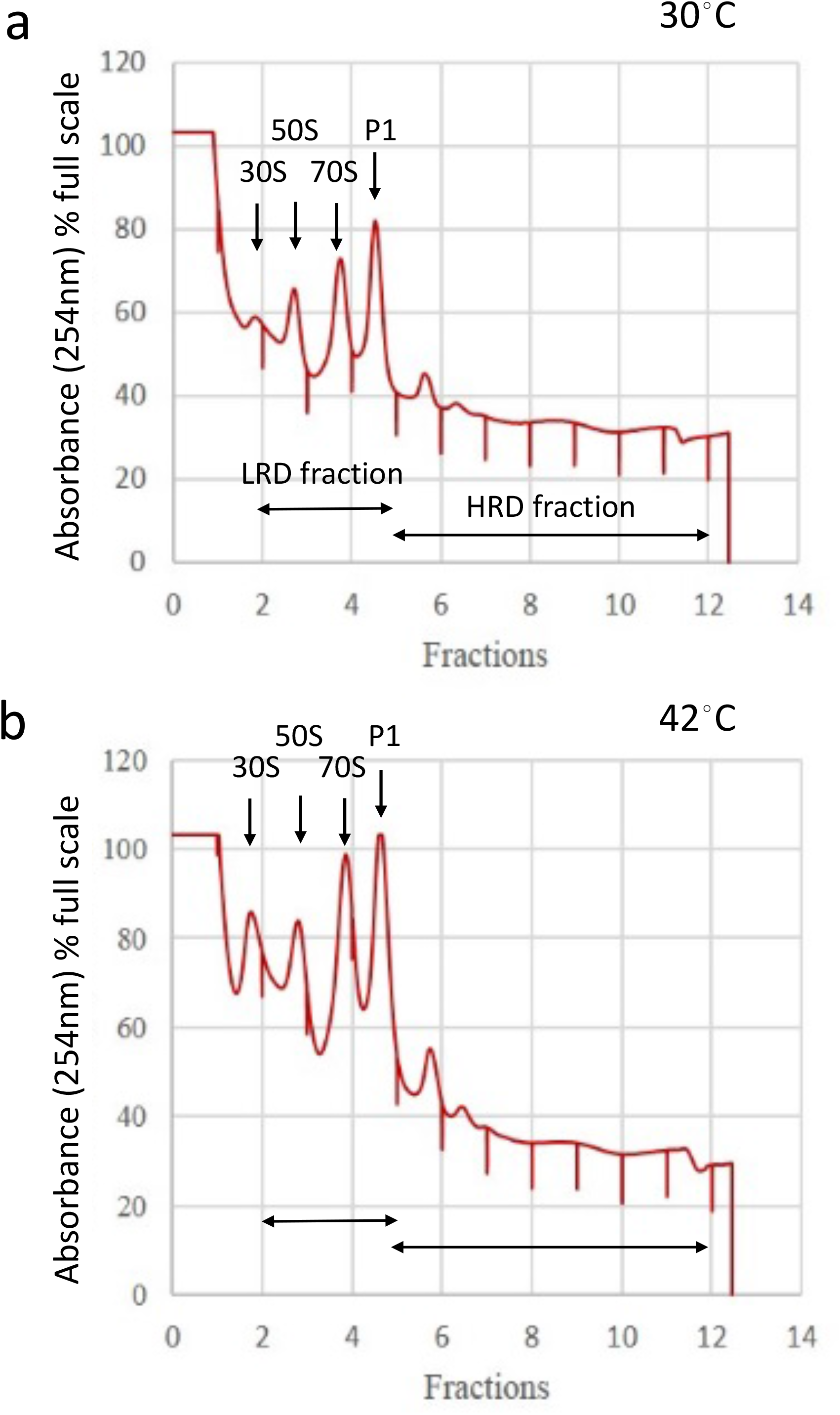
Representative polysome profiles of *S. coelicolor* mycelial extracts from surface grown mycelium cultured at 30°C and heat-shocked at 42°C for 15 mins. Two pools of fractions were studied: the low ribosome density (LRD) pool containing fractions with ribosomal subunits, monosomes and disomes (P1); the high ribosome density (HRD) fraction containing at least 3 ribosomes per mRNA.

In the present study we pooled the low ribosome density fractions (LRD) and the high ribosome density fractions (HRD) separately (Fig. 1) and extracted RNA from the two pooled fractions for subsequent analysis of the differentially ribosome-associated mRNAs.

### Extensive changes in ‘translation efficiency’ occur during heat-shock exposure

Total RNA and polysome-associated RNA fractions were converted into cDNA and analysed on high density whole genome Agilent DNA microarrays using genomic DNA used as a common reference. The microarray data was processed and normalized for differential expression analysis via Rank Products analysis (see Methods). The significantly different gene expression changes at 42°C relative to 30°C, in both transcription and translation were determined and are plotted in Fig. 2 in relation to their genomic location; the data are summarized in **Supplementary Data File 1**. It is striking to observe extensive reduction in transcript level across the genome following heat-shock and, conversely, a more pronounced increase in polysome/monosome associated transcripts following exposure to thermal stress.

**Figure 2.**
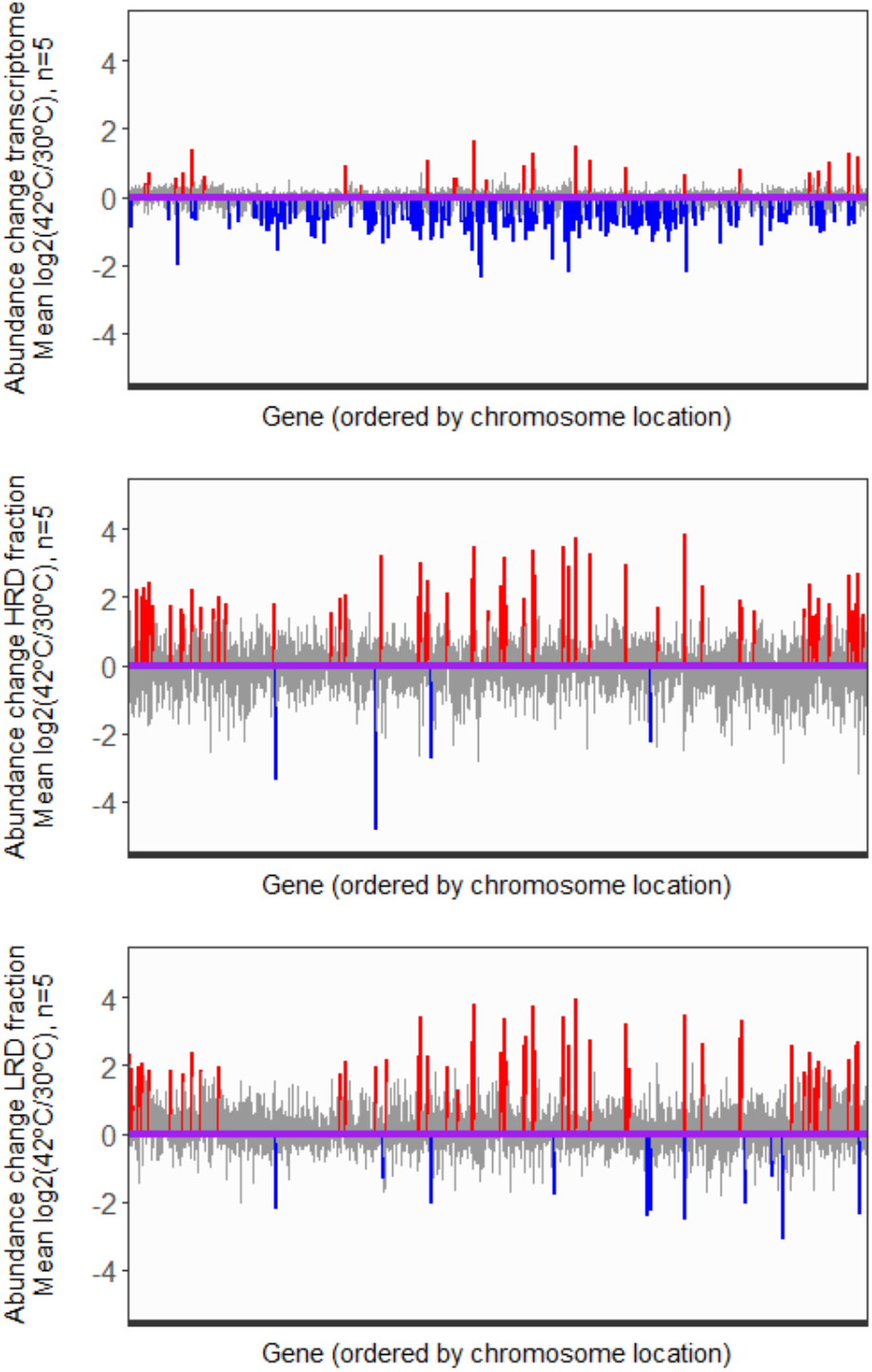
Changes in the average abundance of total mRNA and HRD/LRD associated transcripts following heat shock. Relative log2 fold-change in transcript abundance (average of 5 biological replicates) between 42°C vs 30°C in total mRNA (the transcriptome; top panel), mRNA associated with HRD (middle panel) and mRNA associated with LRD fractions (bottom panel). In each panel, transcripts significantly up-regulated (PFP<=0.1) at the higher temperature are highlighted in red, while those significantly down-regulated (PFP<=0.1) are coloured blue. Non-significant transcripts are coloured light grey, and all transcripts are ordered by their location on the chromosome.

In order to investigate changes in *translational efficiency* (TE), in this case translational induction, the ratio of the transcript abundance in the HRD and LRD fractions to the abundance of the respective transcript in the total RNA fraction, was calculated and compared by Rank Products analysis between the control and heat-shocked samples. In this study, the comparison of ribosome association per transcript is used as a proxy for the efficiency of translation of a gene between two conditions. The results are plotted in Fig 3A-C and suggest that the molecular response to heat shock is governed by the enhancement of translation efficiency of genes over and above any enhancement of transcription.

**Figure 3.**
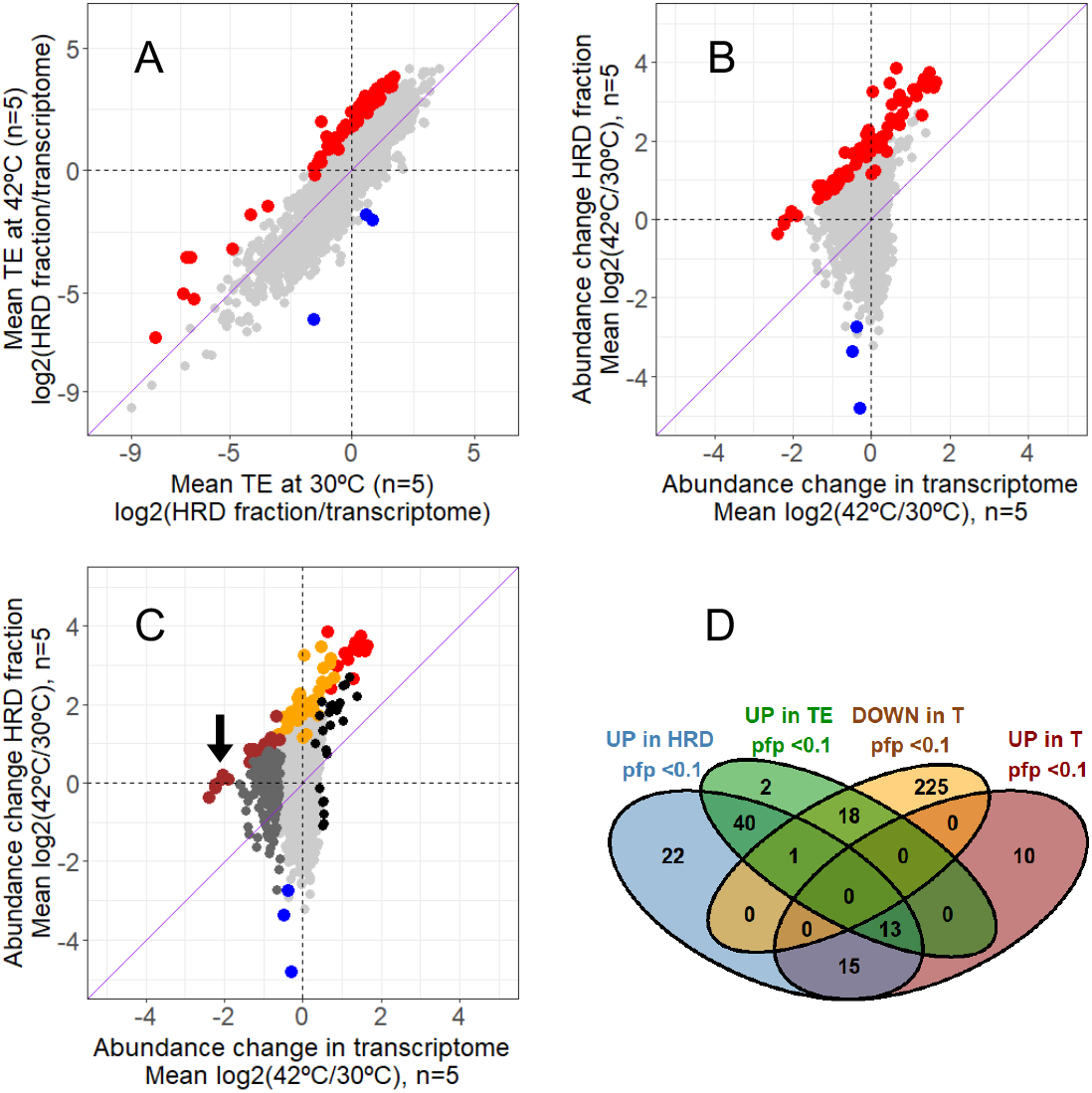
Changes in translational efficiency (TE) are not dependent on transcript abundance (T). A) Identification of transcripts that exhibit a significantly altered translational efficiency (PFP<0.1) following heat shock treatment, based on an analysis of the changes in average abundance of each transcript in the HRD relative to its abundance in the transcriptome. Transcripts that are more efficiently translated at 42°C compared to 30°C are indicated in red, while those showing less efficient translation are shown in blue. Non-significant transcripts are coloured light grey. B) The significant changes in translational efficiency identified in A) (red and blue spots) arise from changes occurring at both the transcriptional and translational level. C) Significant changes in transcript abundance in the transcriptome are not necessarily converted into corresponding changes in HRD association. Black or dark grey spots correspond to transcripts significantly up- or down-regulated (PFP<=0.1), respectively, following heat shock but which exhibit no significant change in abundance in translational efficiency. Transcripts showing significant increases in both transcription and translational efficiency are shown in red, those exhibiting a significant decrease in transcription but with an increased translational efficiency are coloured brown. The arrow indicates the location of the six cold-shock proteins in this group. Orange indicates those transcripts with significantly increased translational efficiency that do not significantly change in abundance in the transcriptome, and blue corresponds to the three transcripts with reduced translational efficiency from A) and B) that do not change significantly in the transcriptome. All nonsignificant transcripts are coloured light grey. Note that the same set of 74 genes is represented in Panel A, B and C. The 4-way Venn diagram in Panel D represents the different subsets of genes that are significantly over-represented in the HRD polysome pool, enhanced translational efficiency (TE), and upregulated or down-regulated at the transcriptional level (T); networks for some of these gene products are illustrated in Figure 4.

The Rank Products analysis described above revealed a total of 74 genes more efficiently translated and only three significantly less translated at 42°C vs 30°C (pfp <0.1); these are shown in Fig 3A and are listed in Table S1.

The respective transcriptional and translational changes of the 74 genes identified as more efficiently translated at 42°C are plotted in Fig. 3B where it can be clearly observed that the changes in translation are of a higher magnitude than changes in transcription. Five groups of transcripts that follow different trends in relation to their abundance changes in polysome association and/or transcription at 42°C vs 30°C are colour-coded differently in Fig. 3C: transcripts which exhibit significant changes in both transcription and translation (red); transcripts which are more efficiently translated but show no significant change in their transcription (orange); transcripts which exhibit a significant decrease in transcription and an increase in translational efficiency (brown); transcripts significantly up or down regulated at the transcriptional level which exibit no significant changes in translational efficiency (black or dark grey); transcripts with reduced translational efficiency (blue). The five different groups of genes highlighted in Fig. 3C reveal the extent of the dynamic response to heat stress which was not fully appreciated in previous studies of *Streptomyces;* our findings contrast with a recent study in *E. coli*, in which a general positive correlation was found between transcription and translation following heat-shock exposure ^28^. Changes in transcription, translation, or both, and translational efficiency following heat-shock are represented in the Venn digram in Fig 3D; an annotated list of the different subsets of genes from the Venn analysis is presented in Table S2.

The picture that emerges from our comprehensive analysis of the heat shock response in *S. coelicolor* is that there is no direct correlation between transcriptional and translational enhancement of gene expression. Instead, we observe a dominant contribution of increased translational efficiency to the heat stress-induced reprogramming of gene expression. A relatively small proportion of genes (13) show both significant transcriptome and translatome-induced changes (Table S2). These genes are designated *‘potentiated’*, a phenomenon that enables rapid enhancement in the production of specific proteins whose roles are essential in dealing with heat-induced cell damage ^15^; in our study all the major heat-shock chaperone and protease-encoding genes are shown to be potentiated, enabling rapid synthesis of key repair and proteolytic processes (Fig. 4; Table S2); the potentiated genes include the DnaK and GroES/GroEL1/GroEL2 molecular chaperone machines and Lon protease.

**Figure 4.**
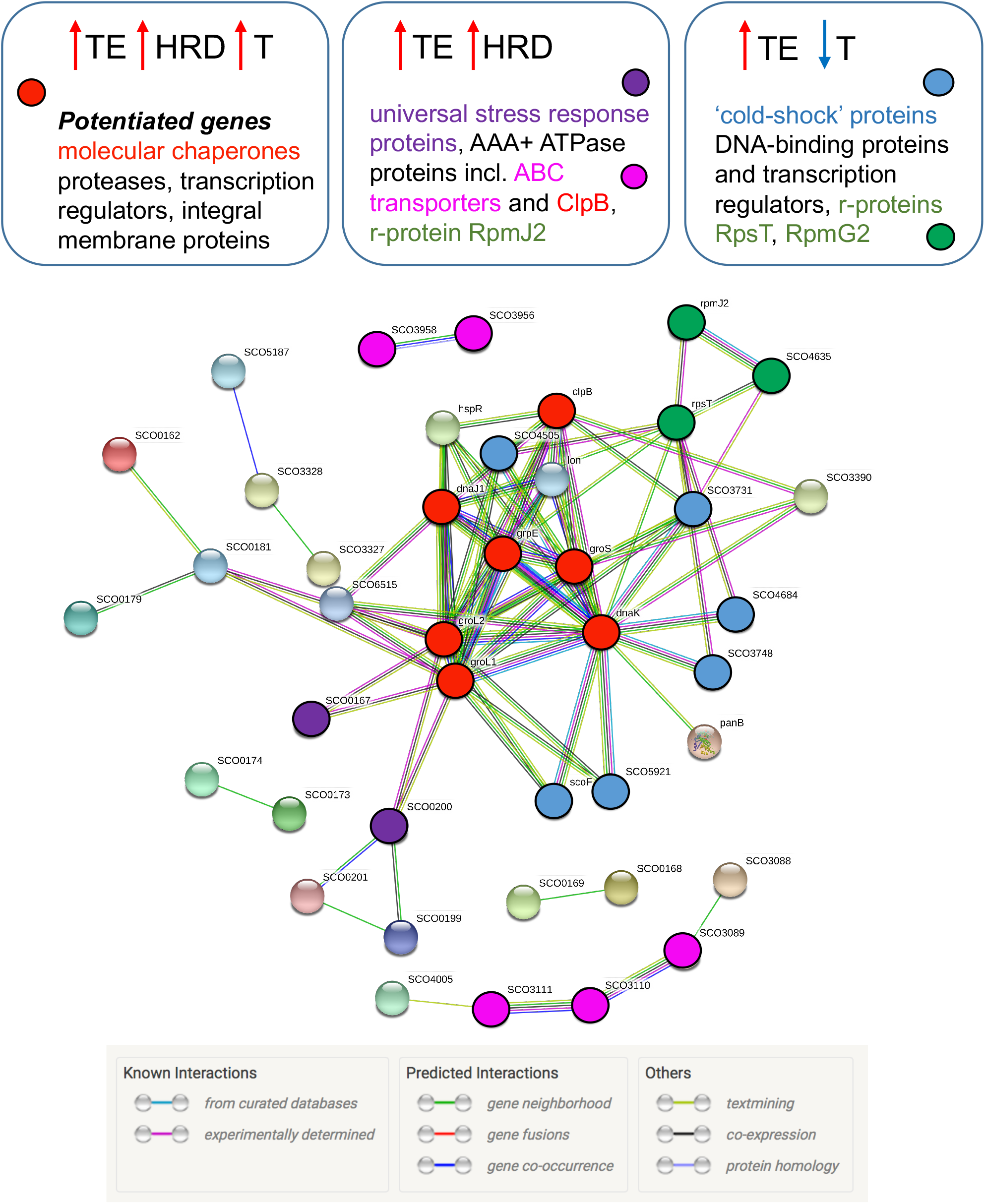
Protein-protein interaction networks from the products of genes with enhanced translational efficiency (TE) following heat-shock. Proteins are shown for which there is a known or predicted interaction, derived from STRING 10.5^44^. Three different groups are highlighted, distinguished by whether transcription (T) is also signicantly altered. Those genes which are ‘potentiated’ (both translationally and transcriptionally induced) include the major molecular chaperone machines. Genes that are translationally induced with no significant change in transcription include universal stress reponse proteins and ABC transporters while genes that are translationally enhanced despite downregulation of transcription include the gene family encoding ‘cold-shock’ proteins. The key protein groups are colour coded and highlighted in boxes above the network.

Functional enrichment analysis of the 74 translationally-induced genes using DAVID software v 6.8 revealed three clusters with enrichment score (E.S.) >1.3 (equivalent to non-log scale of 0.005): (1) Cold-shock protein (CSP)/DNA binding domain (E.S. =6.32); (2) Universal stress protein A (E.S. =2.1) and (3) ABC transporter ATP binding protein (E.S. =1.57). The top scoring Cluster 1 includes genes encoding six ‘cold-shock proteins’ (SCO0527, SCO3731, SCO3748, SCO4505, SCO4684 and SCO5921) and the most enriched GO terms are ‘DNA binding’ and ‘regulation of transcription’; whilst the dominance of CSPs as an enriched cluster might seem counter-intuitive based on their nomenclature, they are also associated with terms such as: ‘DNA binding’, ‘transcription’, ‘transcription regulation’ and stress response, all biological processes that would be required for orchestrating gene expression changes necessary for survival under stress. CSPs are not only required during cold-shock adaptation but also during active growth in *B. subtilis*. ^29^ In fact, SCO0527 and SCO4505 are highly abundant proteins in *S. coelicolor* fermenter cultures grown under nonstress conditions ^30^. CSPs have been proposed to act as RNA chaperones which may facilitate transcription/translation during cold (or heat shock) by preventing the formation of RNA secondary structures ^31^. Cluster 1 also comprises the heat shock transcriptional repressor, HspR, which controls the DnaK chaperone machine, Lon protease and ClpB. Interestingly, two other homologues of HspR, share membership of Cluster 1 (SCO4688 and SCO5917) and, moreover, SCO5917 is the gene most highly associated with polysomes at 42°C relative to 30°C (9.2-fold). Nothing is currently known about the biological roles of SCO4688 and SCO5917 in *Streptomyces*. The remaining three members of Cluster 1 encode the DNA binding proteins, SCO4091, the RNA Polymerase extracytoplasmic sigma factor SCO4005, and SCO3328 *(bdtA)*. SCO4005 encodes a stress-response sigma factor belonging to the *sigE* family, which is also transcriptionally upregulated during the stringent response ^32^ Cluster 2 comprises the Universal Stress Protein, SCO0200, and the hypothetical proteins SCO0167 and SCO0181. It is known that SCO0200 is under the control of the OsdR/OsdS two component system in *S. coelicolor*, the functional orthologue of the DevR/DevS of *Mycobacterium tuberculosis* which controls the dormancy response for surviving the host defence mechanism. The OsdR/OsdS system in *Streptomyces* has evolved a different role in controlling stress and developmental genes to coordinate the onset of morphological development and antibiotic production in response to environmental conditions ^33^. Cluster 3 includes four ATP transporter ATP-binding proteins (AAA+ATPases) also involved in defense mechanisms: SCO3089, SCO3111, SCO3956 and SCO3958. The ClpB (SCO3661) molecular chaperone and Lon protease (SCO5285) are also included in Cluster 3.

It was possible to further subdivide the 74 translationally upregulated genes on the basis of whether their transcription was also significantly influenced following heat-shock (Fig 3D), providing new insights. Three different groups are highlighted in Fig. 4, distinguished by whether transcription was induced, unchanged, or down-regulated. ‘Potentiated’ genes (both translationally and transcriptionally induced) are described above. At least 40 genes are translationally induced but with no significant change in transcription. These include a batch of ‘universal stress response’ proteins and ABC transporters and, notably, 14 of these translationally-induced genes are contained within a 50-gene segment of the ‘redox stress response’ cluster (SCO0162-SCO0212) close to one end of the chromosome^33,34^. These gene products are considered to play a critical role in stress adaptation because the genes are induced only at the level of translation, avoiding the requirement for additional transcription; this regulatory strategy ensures a near instantaneous response to the environmental stressor. By contrast, 19 of the translationally-enhanced genes are significantly down-regulated at the level of transcription. This includes the gene family encoding ‘cold-shock’ proteins (CSPs) and the retention of consistent ribosome occupancy of their mRNAs despite significant reduction in transcript levels suggests an important requirement for a minimal level of production of these proteins. It is possible that the CSPs serve as RNA chaperone proteins, facilitating translation under conditions of thermal stress.

A recent study has established the molecular basis for the interaction of such a cold-shock domain of a CSP with RNA ^35^.

In contrast to the 74 translationally upregulated genes, only three were significantly translationally *repressed* following heat-shock: SCO2629, encoding an integral membrane protein; SCO1580, encoding ArgC, an arginine biosynthetic enzyme, and SCO3229, encoding 4-hydroxymandelic acid synthase, within the gene cluster for the calcium-dependent antibiotic^36^.

### Discriminating features of the genes showing most significant changes in translational efficiency following heat-shock

We next determined whether there were any distinguishing features of the 74 genes that showed the greatest changes in translational efficiency (TE) following heat-shock. Several clear features were found: (a) the most efficiently translated transcripts at 42°C are generally promoter-proximal genes in operons (57 out of 74 are first gene in an operon) and are predominantly leadered transcripts, only one in 74 transcripts is leaderless while across the genome 21% of transcripts are leaderless^14^; (b) The genes encode proteins that are significantly shorter on average than the genome-encoded proteins as a whole; they comprise fewer amino acid residues than would be expected from a random sampling of 1,000 sets of 74 proteins. The value for the significant set is >4 SD away from the mean value for the random sets (Fig. 5A); (c) The amino acid composition of the 74 proteins is significantly different when compared to a random sampling of 1000 sets of 74 proteins (p-value from a one tailed Wilcoxon test) (Fig. 5B); (d) The average G+C content is significantly lower within the 74 genes relative to the rest of the genome. Given an average G+C composition across genome of 72.1%, there is a reduced G+C composition in the 50 bp region upstream of the translation start codon and in the whole coding region of the 74 translationally enhanced genes when compared to a random sampling of 1,000 sets of 74 proteins (p-value from a one tailed Wilcoxon test) (Fig. 5C).

**Figure 5.**
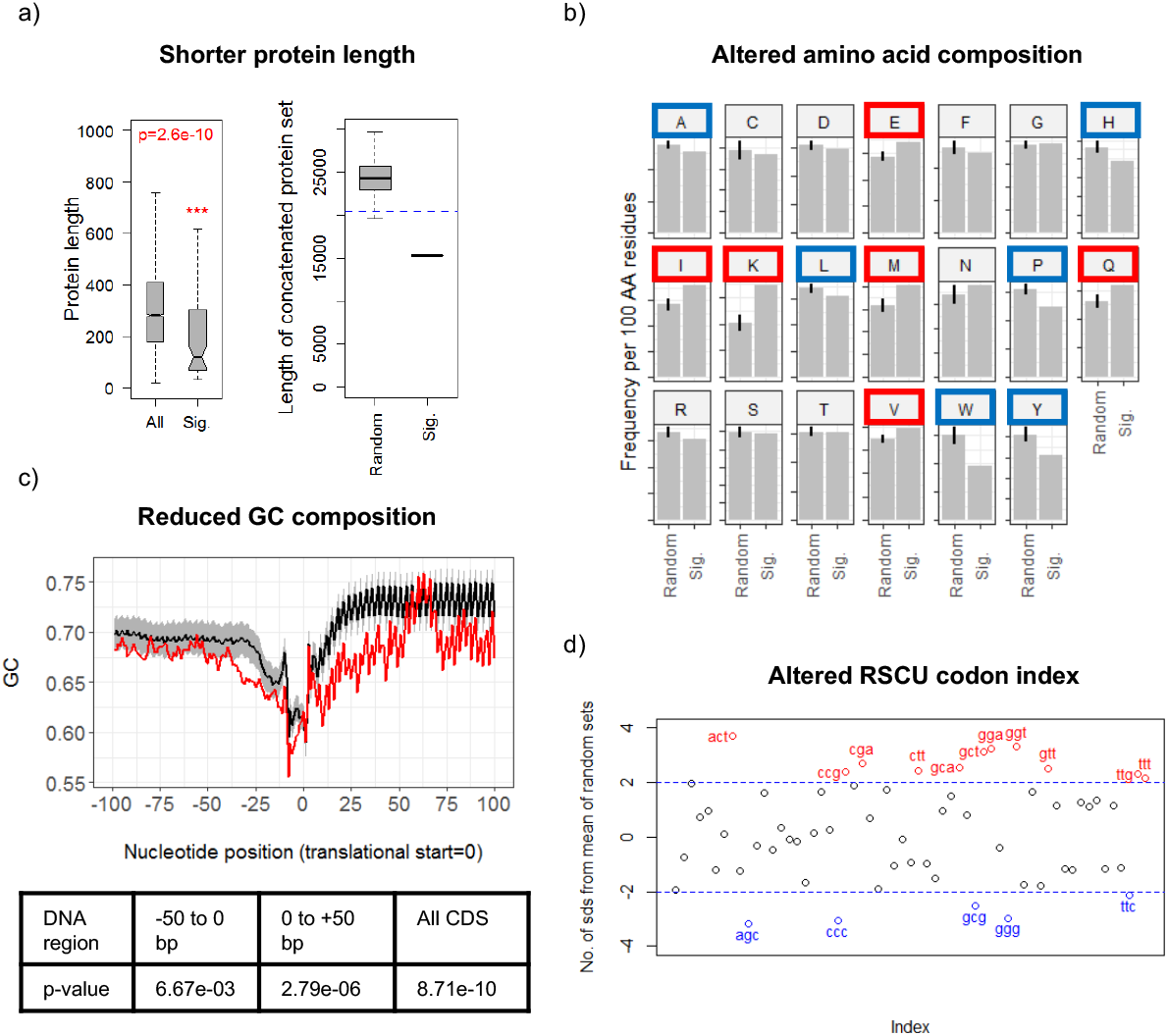
Transcripts that are more efficiently translated at 42°C compared to 30°C are distinguished by their nucleotide content and by the composition of their encoded proteins. (A). Proteins encoded by the more efficiently translated transcripts are significantly shorter than the genome population (left panel, compare Sig. to All; p-value from a one-tailed Wilcoxon test), and when considered as a set contain notably fewer amino acid residues than would be expected from the random sampling of 1000 sets of proteins of equal number (right panel, compare Sig. to Random. The dashed blue line is 2 standard deviations below the mean value for the random sets). The boxes plotted indicate the median, and the upper and lower quartile values of the distributions. B) The amino acid composition of proteins encoded by the more efficiently translated transcripts (Sig.) is different to that observed by random sampling of 1000 sets of proteins of equal number (Random). Amino acids showing an increased frequency of use per 100 residues in the significant set are indicated by red boxes, and blue boxes mark less frequently used amino acids. Frequency values >2 standard deviations higher or lower in the significant set compared to the mean of the random sets are considered to be significantly different. C) The GC nucleotide composition of the more efficiently translated transcripts (upper panel, red line) is reduced compared to that observed by random sampling of 1000 sets of proteins of equal number (upper panel, black line +/−sd (shaded grey)). The average G+C composition across the genome is 72.1%. One-tailed t-tests indicate that the significant set possess significantly reduced G+C composition in the 50 bp region upstream of the translational start site, in the 50 bp of coding sequence downstream of the translational start site, and in the full coding sequence compared with an equivalent analysis of all genes (lower table). D) Relative synonymous codon usage (RSCU) is altered in the genes encoding the more efficiently translated transcripts compared to that observed by random sampling of 1000 sets of proteins of equal number. Codons with RSCU values >2 standard deviations higher (red) or lower (blue) in the significant set compared to the mean of the random sets are considered to be significantly different.

With regard to the amino acid composition (point c, above), some amino acids such as glutamate (E), glutamine (Q), methionine (M), valine (V), lysine (K) and isoleucine (I) are more frequently represented in the proteins while others (alanine (A), histidine (H), leucine (L), proline (P), tryptophan (W), tyrosine (Y)) are less frequently used (Fig. 5B). The finding of elevated glutamate and glutamine content is consistent with our previous report that the HspR transcriptional repressor controls the major E/Q tRNA cluster in addition to the DnaK molecular chaperone machine, *clpB* and *lon*; the increased demand for these tRNAs could be attributable to the increased frequency of these and other amino acids in the identified set of most efficiently translated transcripts at 42°C ^26^. One possible explanation for some of the observed differences in amino acid composition is that it is driven by the lower G+C content of the codons in the translationally enhanced genes (point d above). A good example is the enhanced usage of lysine (encoded by AAn codons) and the reduced usage of arginine (encoded by CGn codons). The apparent increased use of methionine may reflect the increased use of the ATG (methionine) start codon from 61.7% in all *S. coelicolor* genes to 65.7 % in the translationally enahnced genes, compared to the reduction of the GUG (valine) start codon from 35.1% to 30.1%. There is a substantial increase in the rarely used TTG (leucine) start codon from 0.3% to 4.1% in the 74 TE genes.

The bias in synonymous codon usage is also reflected in the choice of stop codons. There is an increase in the use of UAA from 4.4% in all *S. coelicolor* genes to 8.2% in the TE genes, with a co-responding decrease in the use of the UAG codon, 17.4% to 15.0%, and the UGA codon, 78.1% to 76.7%. This may also explain in part the observed reduction in the size of the translationally upregulated proteins (point b above)

### Differences in transcripts identified as differentially translated in the *low ribosome density* (LRD) and *high ribosome density* (HRD/polysome) fractions following heat-shock

Some differences were observed in the distribution of transcripts among the HRD/polysome and LRD pools. Of the 74 genes translationally enhanced at 42°C vs 30°C, 11 transcripts were found to be changed in both the monosome-containing LRD and HRD/polysome pools, while 63 are in the polysome pool only and 4 transcripts are associated with LRD fractions only: SCO0035 encoding an uncharacterized protein of 149 amino acids, SCO4216 encoding an uncharacterized protein of 69 aminoacids, SCO6500 encoding a probable vescicle structural protein of 144 aminoacids and SCO7053, encoding a uncharacterized protein of 139 aminoacids. It has recently been reported that 80S monosomes are translationally active, engaged in translation elongation in *Saccharomyces cerevisiae*. In *S.cerevisiae*, the monosome associated transcripts are generally associated with ORFs smaller than 590 nt and /or have slower initiation rates; they encode predominantly low abundance proteins such as regulatory proteins (kinases and transcription factors)^37^. The subset of transcripts that are preferentially either LRD or polysome associated in our study are illustrated in Fig. 6. It is observed that the number of transcripts partitioning with the LRD fraction increases at 42°C relative to 30°C. Monosome associated transcripts generally encode smaller than average *S.coelicolor* ORFs and are generally derived from poorly expressed genes. The biological significance of the partitioning of some transcripts into the LRD monosome-containing fractions is currently not understood.

**Figure 6.**
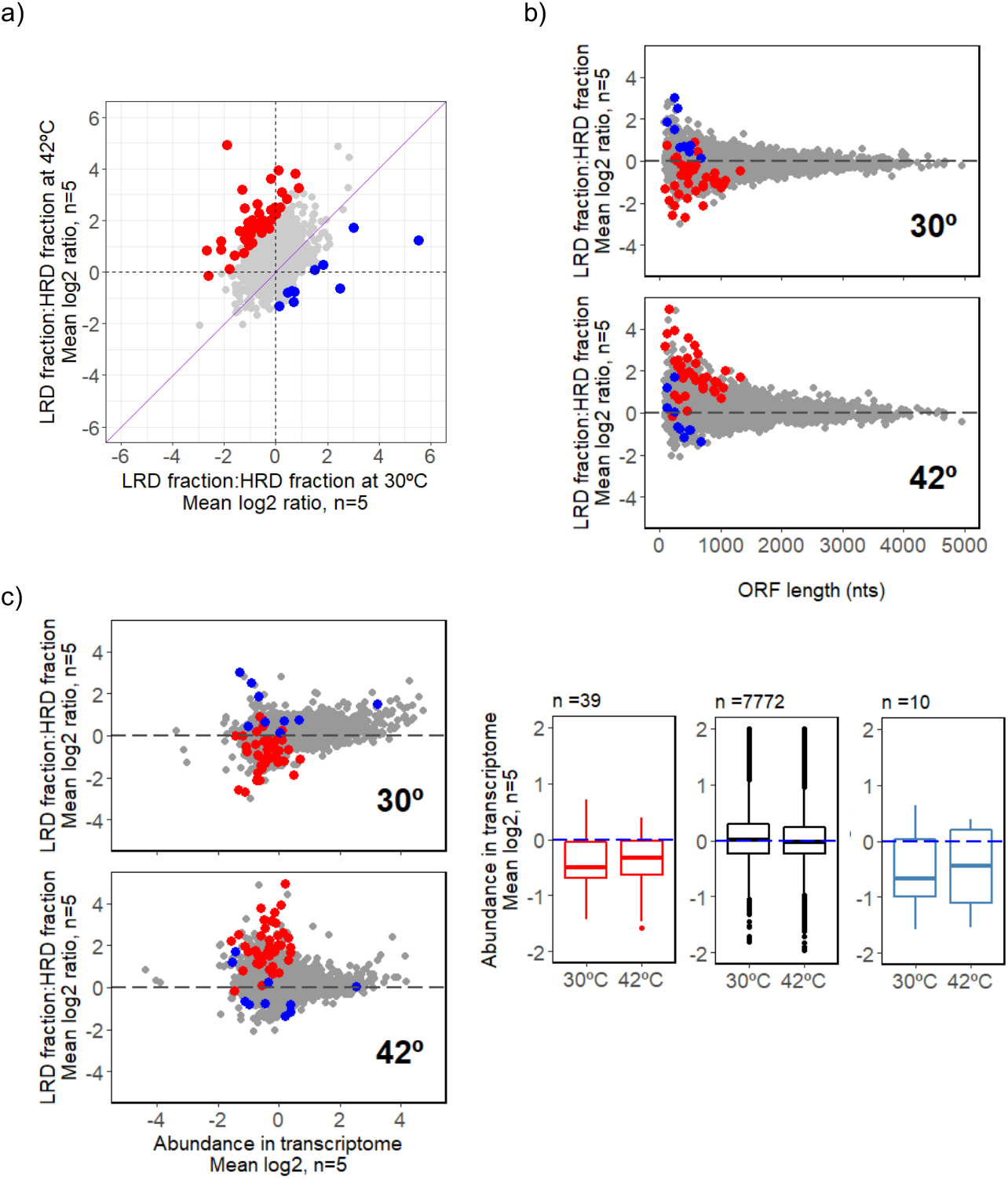
A subset of transcripts show significantly different changes in abundance in the LRD fractions relative to the HRD polysome fractions. a) Identification of transcripts that exhibit a significantly altered partitioning between the LRD and HRD populations (PFP <0.1) following heat-shock treatment, based on an analysis of the changes in abundance of each transcript in the monosome relative to its abundance in the polysome. Transcripts that become significantly more associated with the LRD fraction relative to the the HRD polysome at 42°C compared to 30°C are indicated in red, while those showing a reduced association are shown in blue. Non-significant transcripts are coloured light grey. b) Association of the significant changes identified in a) (red and blue points) with ORF lengths shorter than 1000 nucleotides (nts). c) The significant changes identified in a) (red and blue points) correspond to transcripts with lower than average abundance at both 30°C and 42°C.

### Conclusions

This study has revealed that the predominant upregulatory response to heat-shock in *Streptomyces* is exerted at the translational rather than transcriptional level and has highlighted the likely importance of particular protein families in adaptation to stress that were not previously considered to be part of the heat-shock stimulon. This includes proteins whose expression is only enhanced at the translational level such as a group of ABC transporter systems and a family of cold-shock proteins and specific ribosomal proteins. The role of these newly identified proteins warrants further investigation and their genes provide experimental targets for dissecting the molecular basis for the translational control of the specific mRNAs, the subject of which we know very little about in this important group of antibiotic-producing bacteria.

## METHODS

### Polysome fractionation

S. *coelicolor* MT1110 wild-type spores (2.8 × 10^9^ cfu/ml) were pregerminated in 20 mL of 2YT medium in spring coil-fitted flasks and incubated at 30°C for 8 h in a shaking incubator at 150–200 rpm. The germ tubes were pelleted by centrifugation and re-suspended in 10 mL sterile water. The suspension was mixed by vortexing and the aggregated germ tubes were broken up in an ultrasonic water bath for 10 min. The OD_450_ of 1 ml aliquot was determined and the amount of germ tubes corresponding to 4 × 10^6^ germ tubes per plate was used to inoculate supplemented minimal solid medium (SMMS, Kieser et al., 2000). The inoculated plates were incubated at 30°C up to the desired growth stage conciding with ‘rapid growth phase II’, at the start of red pigmented antibiotic production. At 38–42 h post inoculation, mycelia were heat-shocked at 42°C for 15 min by transferring the cellophane discs onto preequilibrated Whatman paper soaked in water. Following heat-shock, mycelia were rapidly harvested and chloramphenicol treated (80 μg/ml) in order to arrest translation elongation by immersion in chloramphenicol solution. After 1 min incubation at room temperature the cells were chilled on ice and the mycelial pellets were collected by centrifugation at 6,000 rpm/4 min/4°C. The cells were resuspended in 3 ml lysis buffer (20 mM Tris-HCl pH 8.0,140 mM KCl, 1.5 mM MgCl_2_, 0.5 mM DTT, 1% Triton X-100, 80 μg/ml chloramphenicol, 1 mg/ml heparin, protease inhibitor cocktail (Roche) and 100 u/ml RNase out (Thermo Fisher Scientific)). Cells were lysed by mechanical disruption in the presence of glass beads (0.3 cm diameter) in a vortex mixer for 20 sec, four times, with 1 min intervals on ice. The cell extracts were purified by centrifugation at 4,700 rpm/5 min/4°C. The supernatant was transferred to 1.5 ml RNase free tubes and centrifuged again at 9,500 rpm/5 min/4°C. The supernatants were loaded on 10–50% sucrose gradients and ultracentrifuged in a Beckman SW41 rotor at 35,000 rpm / 2 h 40 m/4°C. After centrifugation the sucrose gradients were fractionated in a Teledyne ISCO fractionator (Teledyne Foxy R1 Brandel). The collected 12 × 1 ml fractions were quickly frozen in liquid N2 and stored at −80°C.

### Polysome profiling

The RNA populations associated with monosomes and polysomes were obtained from pooled fractions by affinity purification through mirVANA columns (Life Technologies) which allows for the selective recovery of total RNA (> 17 nt). Nucleic acids from the pooled fractions were bound to filters by repeated centrifugation and disposal of flow-through. The filters were washed twice with 500 μL of wash solution and eluted twice in 50 μL of elution solution preheated to 95°C. The RNA was treated with RNase free DNase (Turbo, Thermo Fisher Scientific) following the recommended manufacturer’s instructions and purified again through RNA clean and concentrator kit TM-5 (ZYMO Research).

Extracted RNA was QC-checked using an Agilent BioAnalyzer 2100, Cy-dye labelled and hybridized separately onto high density whole genome *Streptomyces coelicolor* microarrays (2 × 105K probe-set) using genomic DNA as common reference as described in ^38^.

### Transcriptome analysis

Mycelia from the same bacterial cultures used in the polysome profiling experiments were treated with RNA Protect Bacteria Reagent and total RNA was extracted following a modified version of the method as described in ^34^ In brief, the mycelia was harvest from four plates with a spatula and, after treatment with RNA protect Bacteria Reagent (Qiagen), was resuspended and lysed in 200 μL Tris-EDTA plus Lysozyme (15mg/ml). Then 600 μL of RPE buffer from the Qiagen RNeasy mini kit was added. The mixture was transferred to a sterile RNase free 2 mL micro centrifuge tube with a single stainless steel bead. The tube was placed in the tissue lyser (Qiagen) and shaken vigorously for 2 min, rotated 180° and shaken for another 2 min. The samples were centrifuged for 2 min and the supernatant was transferred to a fresh tube, phenol/chloroform extracted and chloroform extracted. The aqueous phase was passed through a gDNA eliminator column from the RNeasy mini Plus kit (Qiagen). The RNA was purified from the lysate using the mirVana kit columns (Thermo Fisher Scientific) and following the steps recommended by the manufacturer until elution in RNase free water.

The total RNA and RNA from LRD- and HRD-associated fractions were quantified using a Nanodrop ND-2000 Spectrophotometer (Labtech) and Quality control checked with the Agilent Bioanalyser. All RNA had a R.I.N (RNA Integrity Number) >7 before proceeding to RNA labeling and hybridization.

A total of five biological replicates were processed for transcriptome and translatome analysis. Each RNA sample (amount ranging from 2 to 10 μg depending on the availability) was labelled with Cy3dCTP using the method described in ^34^

Each cDNA labelled with Cy3 dCTP was cohybridized with genomic DNA used as common reference labelled with Cy5-dCTP onto high density whole genome *Streptomyces coelicolor* microarrays (2 ×105K probe-set)^38^. The microarray data has been assigned ArrayExpress accession number E-MTAB-6206.

## Data Analysis

Agilent Feature extraction files were read into R ^39^ using the limma package ^40^, and used to produce a final gene expression matrix (GEM) comprising the log_2_(signal)-log_2_(control) values, where signal and control are the “gProcessedSignal” and “rProcessedSignal” measurements, repectively. Log2 ratios involving negative signal values or negative control values were replaced with the lowest non-negative values of the distribution. Cases with negative values in both signal and control were assigned a Log2 ratio value of 0. Each column of the GEM (each microarray) was median-centred followed by scale normalisation across arrays (columns), as defined by the ‘withinArray’ and ‘BetweenArray’ normalisation functions in the limma R package ^40^. The between array normalisation was performed in two batches: one for the total RNA samples and a second batch for the HRD and LRD-associated RNA samples since the distribution of values for the polysome- and monosome-associated RNAs was found to be different from that of the total RNA samples. After normalization, control probes were filtered out (remaining probes n= 103,694), then data from probes targeting the coding region of the same gene were averaged to produce a single transcript abundance value per gene (remaining features n= 7,821). These values were used to calculate translational efficiency (TE) values for each transcript, defined as the ratio of the transcript abundance in the HRD fraction to the abundance of the transcript in the total RNA fraction. Rank products analysis ^41^ was used to test for differential transcript abundance or TE/ratios of transcript abundance between conditions using a percent false positive (pfp) significance cut off of <=0.1. Analyses of the G+C base composition of DNA sequences, the codon usage preference of open reading frames and the amino acid composition of encoded proteins were performed using the seqinr R package ^42^. Where indicated in the text, random sampling and t-test or wilcox significance testing were performed using R base functions. Random sampling of genes from the genome considered all protein-encoding genes excluding pseudogenes.

## Functional enrichment analysis

Functional enrichment analysis was carried out using DAVID software v 6.8 (all *S. coelicolor* genes as background, medium setting for classification stringency, enrichment P value for individual sets of genes (ease score) set at 0.05) ^43^ and STRING v 10.5 protein association network ^44^ with default settings.

## Acknowledgements

The authors are grateful to Andre Gerber for technical advice and troubleshooting the initial polysome profiling experiments in *Streptomyces*, Dr Belinda Hall and Dr Rachel Simmonds for technical help with the ISCO gradient fractionator. We also thank Andre Gerber and Nicolas Locker for helpful discussions.

## Authors’ contributions

G.B., E.E.L., G.R.S. and C.P.S. conceived the study, G.B. and R.P. performed the experiments, E.E.L., A.H., C.M.L., G.B., G.R.S., and C.P.S. and analysed and interpreted the data and G.B., A.H., G.R.S. and C.P.S. wrote the manuscript.

## Supplementary files

### Supplementary Data File 1. (format: .xlsx)

Normalised microarray data and Rank Products analysis of the transcriptome and translatome data reported in this study. The first worksheet, README, details the contents of the data file

### Table S1 (format: xlsx)

Significant translationally enhanced transcripts at 42°C >30°C (74 genes) and 30°C >42°C (3 genes). List of genes (column A); abundance fold changes at 42°C/30°C (74 genes) and 30°C/42°C (3 genes) (column B), probability of false prediction from rank product analysis (column C), significantly different also at transcription at 42°C>30°C where 0 denotes not significant, 1 significantly upregulated and −1 significantly downregulated, protein identifier (column F) protein name (column G) gene name (column H) gene length (column I) gene ontology (columns L-N) and gene name (column R)

### Table S2 (format: xlsx)

Gene lists represented in the Venn Diagram (Fig 3D)

### Legend

Gene = SCO number of gene

Product = predicted protein product

Symbol = gene name symbol

Protein.Length = length in amino acid residues of the predicted protein

Significance_LRD = significance call from rank products analysis of transcript abundance changes in the LRD fraction following heat-shock: 0=not significant, 1=significantly increased at 42°C, −1 = significantly decreased at 42°C

Significance_HRD = significance call from rank products analysis of transcript abundance changes in the HRD fraction following heat-shock: 0=not significant, 1=significantly increased at 42°C, −1 = significantly decreased at 42°C

Significance_T = significance call from rank products analysis of transcript abundance changes in the transcriptome following heat-shock: 0=not significant, 1=significantly increased at 42°C, −1 = significantly decreased at 42°C

Significance_TE = significance call from rank products analysis of changes in TE following heat-shock: 0 = not significant, 1= significantly increased at 42°C, −1 = significantly decreased at 42°C

Significance_LRDvT = significance call from rank products analysis of changes in ratio of LRD:T following heat-shock: 0=not significant, 1=significantly increased at 42°C, −1 = significantly decreased at 42°C

Significance_LRDvHRD = significance call from rank products analysis of changes in ratio of LRD:HRD following heat-shock: 0=not significant, 1=significantly increased at 42°C, −1 = significantly decreased at 42°C.

### Table S3 (format: docx)

Start codon and stop codon preferences in the 73 TE genes (note that one of the 74 original TE UP genes is a pseudogene)

